# Effect of foot position during plantarflexion on the neural drive to the *gastrocnemii* in runners with Achilles tendinopathy

**DOI:** 10.1101/2023.08.14.553177

**Authors:** Gabriel L. Fernandes, Lucas B. R. Orssatto, Anthony J. Shield, Gabriel S. Trajano

## Abstract

Runners with Achilles tendinopathy (AT) have reduced neural drive to the *gastrocnemius lateralis* (GL). This study investigated if the strategy of pointing feet-inward (*feet-in*) during isometric plantarflexion would increase *gastrocnemius lateralis* electromyography root mean square amplitude (RMS) and motor unit discharge rates, compared to feet-in neutral position (*feet-neutral*), in runners with Achilles Tendinopathy (AT). High-density electromyograms were recorded from *gastrocnemius lateralis* and *medialis*, during 20-s *feet-in* and *feet-neutral* isometric heel raise, in runners with (n=18) and without (n=19) AT. During *feet-in*, GL RMS was higher during *feet-in* in both groups and GM RMS was lower only during *feet-in* in the AT. Conversely, motor unit discharge rates were lower during *feet-in* in GL (p<0.001) and in GM in the AT group. The AT group had lower *triceps surae* endurance during single leg heel raise. In summary, *feet-in* increases GL RMS in both groups, conversely reducing motor unit discharge rates in the AT group, compared to *feet-neutral*. Additionally, *feet-in* reduces GM RMS and motor unit discharge rates only in the AT group, compared to *feet-neutral*. This would shift the *gastrocnemius lateralis/medialis* ratio excitation, favouring *gastrocnemius lateralis*. Nonetheless, while this strategy holds promise, it remains uncertain whether performing plantarflexion exercise with feet pointed inwards would provide additional benefits for the treatment of runners with Achilles tendinopathy. Our findings suggest that the increased GL RMS during *feet-in* is effective in increasing GL excitation but not as consequence of increased MUDR and, but it might be a result of recruitment of more motor units.

## Introduction

Achilles tendinopathy, is a highly prevalent injury among runners (Lagas et al., 2020; Mousavi et al., 2019) and is associated with deficits in *triceps surae* performance (Sancho et al., 2019; Silbernagel et al., 2006). Runners with acute presentations of Achilles tendinopathy (<3 months) exhibit lower neural drive to *gastrocnemius lateralis* during submaximal triceps surae isometric force (Crouzier et al., 2020) and dynamic tasks, such as bilateral and unilateral heel drops (Mylle et al., 2023). Lower neural drive to *gastrocnemius lateralis* is also present in chronic (> 3 months) Achilles tendinopathy (Fernandes et al., 2022), evidenced by lower motor unit discharge rate during submaximal isometric contraction (Fernandes et al., 2022). This behaviour seems to be specific to *gastrocnemius lateralis*; since there seems to be no differences in neural drive to *gastrocnemius medialis* and *soleus* compared to healthy controls (Crouzier et al., 2020; Fernandes et al., 2022; Mylle et al., 2023). Individuals with Achilles tendinopathy seem to have a compensatory *triceps surae* muscle coordination strategy, favouring *soleus* over *gastrocnemius lateralis* (Mylle et al., 2023), which reinforces the idea of impaired *gastrocnemius lateralis* neural drive. These deficits in *gastrocnemius lateralis* neural drive observed in runners with Achilles tendinopathy (Crouzier et al., 2020; Fernandes et al., 2022) may negatively impact Achilles intratendinous load distribution; increasing tendon strain and potentially contributing to tendinopathy (Bojsen-Møller et al., 2004; Cook & Purdam, 2009; Hug & Tucker, 2017). Developing rehabilitation strategies that aim to increase *gastrocnemius lateralis* activation and contribution to *triceps surae* force could be a promising approach to improve plantar flexor neuromuscular control and provide better rehabilitation outcomes for this population.

Although a synergist muscle group during ankle plantarflexion, motor units from individual muscles of the *triceps surae* have partially independent neural drive (Hug et al., 2020). Changing the orientation of the foot inwards or outwards during plantarflexion can increase the root mean square amplitude (RMS) and motor unit discharge rates to either *gastrocnemius lateralis* or *medialis*, respectively (Crouzier et al., 2022). Specifically, plantarflexion exercise performed with the feet pointed inwards increases global *gastrocnemius lateralis* sEMG amplitude (Crouzier et al., 2024) and motor unit discharge rates (Hug et al., 2020) in healthy individuals. This is further supported by the greater *gastrocnemius lateralis* hypertrophy observed after resistance training performed with feet pointed inwards (Nunes et al., 2020). Whereas positioning the feet outwards has the same affect to *gastrocnemius medialis*. Thus, performing rehabilitation exercises with feet pointing inwards may be beneficial to increase *gastrocnemius lateralis* contribution to plantarflexion in healthy individuals. However, it remains unclear if this assumption is reproducible in runners with chronic Achille tendinopathy, a population with reduced neural drive to *gastrocnemius lateralis*.

This study aimed to investigate if the inwards foot position would increase the neural drive to *gastrocnemius lateralis* in runners with Achilles tendinopathy compared to the neutral position. Participants performed bilateral standing isometric plantarflexion contractions with *feet-in* and *feet-neutral.* We hypothesised that the feet pointed inwards (*feet-in*) condition would: i) increase EMG signal amplitude and motor unit mean discharge rates of *gastrocnemius lateralis* in both Achilles tendinopathy and controls; and ii) reduce motor unit mean discharge rate and RMS of *gastrocnemius medialis* in both groups, compared to *feetneutral*. These data could guide the development of novel rehabilitation interventions, using different foot position during ankle plantarflexion, to increase neural drive to *gastrocnemius lateralis* in runners with chronic Achilles tendinopathy.

## Methods

### Participants and ethical considerations

Thirty-seven endurance runners were recruited for this study, 18 runners with Achilles tendinopathy and 19 controls (characteristics reported in **TABLE 1**). Volunteers were recruited from local running clubs around Southeast Queensland, Australia, via email and social media. Diagnosis of Achilles tendinopathy was confirmed by a senior physiotherapist during examination, on the basis that patients presented with localised Achilles tendon pain for more than three months, pain was provoked by physical activities in a dose dependent way and by palpation of the tendon. Average symptom presentation of Achilles tendinopathy was 4.9 ± 6.3 years. Volunteers were excluded if they had previous rupture of or surgery to the Achilles tendon; clinical findings indicating a differential diagnosis for the Achilles tendon pain (such as tendon sheath inflammation, an acute tendon tear, plantaris or tibialis posterior tendon involvement, neural involvement); VISA-A score > 90 points for Achilles tendinopathy group and < 100 for the control group; any other musculoskeletal injuries of the lower limb of the affected leg; any neurological disorder; or mental health issues affecting consent. All participants were free of cardiac, pulmonary, renal or endocrine (Knobloch, 2016) comorbidities. Prior to testing, all participants read and signed a detailed informed consent document and completed the VISA-A questionnaire (Martin et al., 2018). The Achilles tendinopathy group also completed the Tampa scale of Kinesiophobia to identify if kinesiophobia was a factor influencing *triceps surae* performance (Chimenti et al., 2021; Silbernagel et al., 2011). *Triceps surae* endurance deficits are widely reported in runners with Achilles tendinopathy (Fernandes et al., 2021; O’Neill et al., 2019; Quarmby et al., 2023; Silbernagel et al., 2007). Thus, a single leg heel raise endurance test (Hébert-Losier et al., 2017; Id et al., 2021) was used as measure of *triceps surae* performance to help characterise groups and compare changes in *gastrocnemii* motor unit discharge rate and RMS related to foot position. This study was approved by the Queensland University of Technology Human Research and Ethics Committee in line with the Declaration of Helsinki. All participants signed an informed consent prior participation in the study.

**TABLE 1.** Participant characteristics.

| Variable | Achilles tendinopathy (n=18) | Control (n=19) | Mean difference |
| --- | --- | --- | --- |
| Age (years) | 44.2 ± 8.9 | 35.4 ± 6.5 | 8.8 ± 6.2 |
| Sex | 12 males (66%) | 10 males (53%) | — |
| Height (cm) | 165.5 ± 38.5 | 172.1 ± 11.3 | -6.6 ± 4.7 |
| Body Mass (Kg) | 71.5 ± 12.8 | 67.5 ± 13.8 | 4.0 ± 2.8 |
| Running mileage<br>km/week | 43.6 ± 26.8 | 42.2 ± 29.5 | 1.4 (± 1.0) |
| Running experience | 18.8 ± 13.3 | 12.6 ± 8.5 | 6.2 ± 4.4 |
| VISA-A score | 65.5 ± 13.2 | 100 (0) | -34.5 ± 24.4 |

Data represent mean and ± standard deviation (SD).

#### Study design

As part of the inclusion criteria, participants were requested to complete a VISA-A questionnaire and send it via email before their scheduled lab visit. Additionally, a Tampa scale of Kinesiophobia questionnaire was sent via email prior to the visit. Testing involved a single laboratory visit, in which participants performed a warm-up followed by maximal plantarflexor isometric contractions in an isokinetic dynamometer. Thereafter, participants performed four 20-s bilateral standing plantarflexion isometric contractions, two with feet positioned in neutral (*feet-neutral)* and two with feet-inward (*feet-in).* Lastly, they performed a unilateral heel raises to failure (SLHR) using their affected/dominant leg for the Achilles and controls respectively.

#### Data collection procedures

Plantarflexor isometric peak torque was measured using an isokinetic dynamometer (Biodex Medical Systems, Shirley, New York). For the bilateral Achilles tendinopathy presentations (n=9), the most symptomatic leg was used and for the control group, the dominant leg was used for testing. Participants were seated (75 degrees of hip flexion) with their knee straight and with the ankle joint at 10 degrees of plantarflexion. Warm up consisted of 2 × 4 s isometric contractions of each participant’s perceived 30, 50, 70 and 80% maximal voluntary isometric contraction intensity. After warm-up, participants performed at least 3 maximal voluntary isometric contractions with 2 minutes rest between attempts, until <5% variation was observed between contractions. No more than 4 attempts were performed. The attempt with the highest torque was analysed. Participants then performed, 8 bilateral standing heel-raises, 4 in each condition, *feet-neutral* and *feet-in*; in a randomised order with 1 minute rest between contractions. *Feet-neutral* condition was considered to be each participant’s comfortable stance with feet pointing forward; whereas for *feet-in* condition, participants internally rotated their hips so that the long axis of their feet were at 90-degrees to each other (Hug et al., 2020). All participants were able to achieve and comfortably stay in the *feet-in* position for testing. They were instructed to perform a heel raise to full range followed by a slow drop until their heels lightly touched a 6.5 cm height soft foam (supervised by tester) and to maintain their position for isometric testing. Additionally, participants were instructed to lightly touch a table positioned in front of them with their hands to minimise oscillation and aid in maintaining balance. Once in position, a 20 second isometric contraction was recorded. After all contractions were finalised, participants were given ∼10 min rest, seated in a chair. Thereafter, a single leg heel raise (SLHR) endurance test was performed, using a metronome set to 60 bpm (Hébert-Losier et al., 2017; Id et al., 2021) to control tempo (1 second up, 1 second down). Participants were instructed to go as high as possible and come down to fully touch their heel on the ground before going up again. Test was ceased once performance was no longer correct for 2 consecutive repetitions (as indicated by reduced plantarflexion range, bending knees, hip hinge, or other compensatory strategies or the inability to maintain the pace).

#### High-density electromyograms recordings and analyses

HD-EMG (Sessantaquattro, OTBioelettronica, Torino, Italy) signals were recorded from *gastrocnemius medialis* and *gastrocnemius lateralis* using OT Biolab+ software (version 1.3.0., OTBioelettronica, Torino, Italy) during two isometric plantarflexion contractions per condition and muscle. A b-mode ultrasound (ESAOTE, MyLabSeven. Genova, Italy) was used to identify muscle fibre orientation and muscle boundaries, to guide electrode positioning. To ensure good skin-electrode contact and conductance, a bi-adhesive layer with a conductive paste was used. One 64-channel electrode grid (GR08MM1305, OTBioelettronica, Torino, Italy) was placed on *gastrocnemius medialis* and another one on *gastrocnemius lateralis*. The ground strap electrode (WS2, OTBioelettronica, Torino, Italy) was dampened and positioned around the ankle joint of the tested leg. EMG signals were recorded in monopolar mode, amplified (256x), band-passed filtered (10-500Hz) and converted to digital signal at 2048Hz by a 16-bit wireless amplifier (Sessantaquattro, OTBioelettronica, Torino, Italy), before being stored for offline analysis. Although using 2×32 channels (one electrode for each muscle) would have enabled measures of both muscles during the same contraction, a 32-channel electrode yielded a lower number of motor units, in particular for *gastrocnemius lateralis*. Additionally, the larger 64-channel electrode offered a greater surface area, providing more comprehensive coverage of the muscle’s electrical activity. HD-EMG signals were recorded and analysed offline, decomposed into motor unit spike trains with specialised software using blind source separation decomposition technique (DEMUSE, v.4.1; The University of Maribor, Slovenia) (Del Vecchio et al., 2020). Motor units were tracked across all four contractions (two for each condition) and muscles. Tracking motor units across conditions allows a better understanding of the effect of foot position on the discharge patterns of the same motor unit and therefore, the influence of each condition on the neural drive to each muscle. All contractions were analysed, with motor units subjected to visual inspection by a skilled operator to identify any erroneous discharge times. Manual inspection is required to reduce automatic decomposition discharge errors and improve data reliability (Del Vecchio et al., 2020; Martinez-Valdes et al., 2016). Only motor units with a pulse-to-noise ratio (PNR) >30dB, sensitivity > 90%, were used for data analysis (Del Vecchio et al., 2019, 2020). Mean motor unit discharge rates data were analysed using *openhdemg* opensource software (Valli et al., 2024).

#### Root-mean-square electromyography envelope

The differential values of EMG data recorded from the HD-EMG electrodes were analysed longitudinally in the direction of the muscle fibre between the subsequent electrodes, then the EMG root-mean-square (RMS) envelope was calculated. RMS values of all bipolar EMG signals were calculated over epochs of 0.5 s. Thereafter, each RMS channel was normalised (RMS) as a percentage of their respective peak RMS obtained during maximal voluntary isometric contractions, for each condition, also calculated over a 0.5 s period around the maximal values for each bipolar channel (Ema et al., 2020). EMG RMS was calculated for the two contractions and conditions (*feet-in* and *feet-neutral*) for each muscle. Channels were longitudinally classified into proximal (channels 1-4), intermediate (5-8) and distal (9-12) clusters (Rainoldi et al., 2004) to explore potential regional dominance of activation. EMG normalised RMS is referred from now on as RMS.

#### Statistical analysis

All analyses were performed using R studio (version 2023.09.01+494). Linear mixed-effect models were fitted using the *lme4 and lmerTest* (Bates et al., 2015) packages. Linear mixed-effect model using participant ID as random intercept, were used to compare all channels from RMS from *gastrocnemius medialis* and *gastrocnemius lateralis* between conditions (*feet-neutral* and *feet-in*) and groups. A potential effect of cluster and contraction were explored by including them as random effects in the model. Since no effect was observed, cluster and contraction were then removed from the models for the final analysis. Finally, mean RMS from all values obtained from the intermediate cluster comprising 20 channels was chosen for analysis. This decision was based on their location at the centre of the muscle belly, which is widely recognized as the preferred sensor site for sEMG recordings (Rainoldi et al., 2004; Zaheer et al., 2012). Additionally, separate linear mixed-effect models were used to compare motor unit discharge rates from tracked motor units from *gastrocnemius medialis* and *gastrocnemius lateralis* between conditions (*feet-neutral* and *feet-in*) and groups (Achilles tendinopathy and control). The models were tested using participant ID as random intercept to account for the correlation between repeated observations for each participant. Motor unit ID was also included in the model as random intercept to estimate individual change in neural drive in the same motor unit, influenced by experimental condition. Finally, a model testing motor unit discharge rates using participant ID and motor unit ID as random intercepts was used. The final model was selected from a series of candidate models, based on the smallest Bayesian information criteria (BIC) and Akaike information criterion (AIC) values. Additionally, an analysis of covariance (ANCOVA) was used to compare peak isometric torque between groups, using body mass as covariate and a Welch two sample t-test was use for single leg heel raise (SLHR) analysis. The estimated marginal mean difference and 95% confidence intervals (CI) for motor unit mean discharge rate between groups, were determined using the *emmeans* package (Lenth, 2016). Normality assumptions were confirmed by analysis of the histogram of residuals, Q-Q Plot and the residual-predicted scatterplot, and Shapiro-Wilk normality test. *QQnorm* function in R produces a qqplot (quantile-quantile plot) and a theoretical qqline to identify potential outliers based on values below and above 0.25 and 0.75 quartiles of the residuals. When required, a sensitivity test was utilised to assess the influence of outliers identified by the models in the output, and these are reported in the results section. When a significant effect was observed pairwise comparisons were used. Statistical significance was accepted at P ≤ 0.05 for all tests. Omega squared (ω2) effect sizes derived from the F ratios is presented for each linear mixed models analyses (0–0.01, very small; 0.01–0.06, small; 0.06–0.14, moderate; and >0.14, large) (Lakens et al., 2013). Cohen’s *d* effect size is presented as “*d”* for the t-test. Data is presented as mean ± SD or mean (95% confidence interval lower to upper limits).

## Results

### Normalised RMS

*Gastrocnemius lateralis* RMS was different between condition (F=58.11, p<0.001, ω2=0.02), but not groups (F=0.03, p=0.85), nor did it show a condition × group interaction (F=0.69, p=0.40). Mean difference for RMS was 3.4 (2.5 to 4.2) % higher during *feet-in* (**FIGURE 1A**). *Gastrocnemius medialis* RMS analysis showed a condition × group interaction (F= 9.25, p=0.02, ω2<0.001). RMS was 3.4 (-5.4 to -1.4) % lower during *feet-in*, in the Achilles tendinopathy group and no different in controls 0.2 (-1.9 to 1.6) % (**FIGURE 1B**).

**FIGURE 1.**
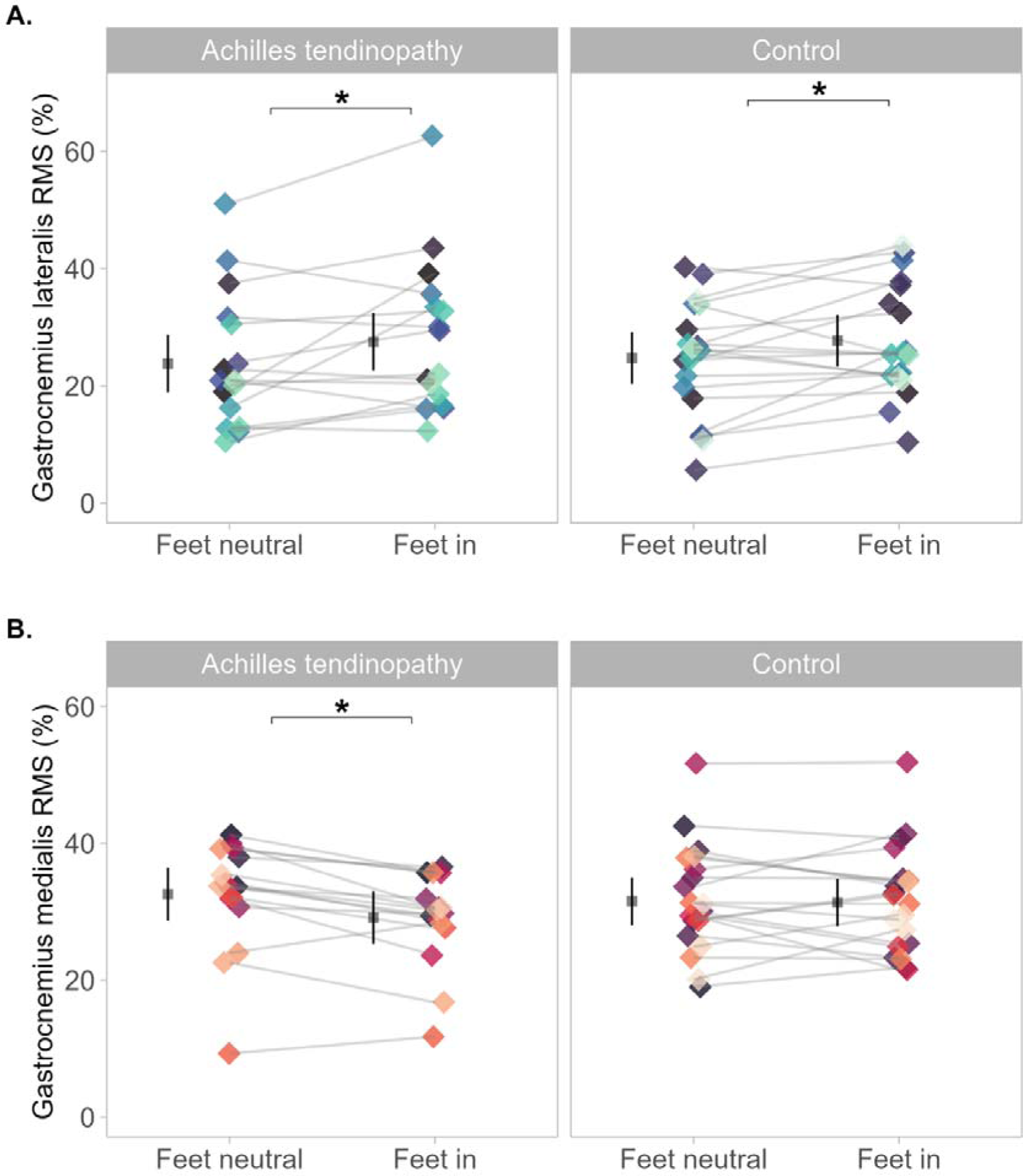
Mean normalised root mean squared (RMS) with *feet-neutral* and *feet-in* between groups. Normalised RMS represents a % of peak RMS. Each dot represents individual mean normalised RMS across condition, coloured by participants. **A.** represents data from *gastrocnemius lateralis* and shows higher RMS during *feet-in* (p< 0.001) in both groups. **B.** displays data from *gastrocnemius medialis*; and shows lower RMS during *feet-in* (p=0.002) only in the Achilles tendinopathy group. Mean and 95% confidence interval are offset to the left to facilitate visualisation. * Denotes statistical difference.

### Motor unit analysis

We found a total of 785 motor units tracked across contractions and conditions. 428 motor units in the Achilles tendinopathy group (11.8 ± 11.2 per participant) and 357 motor units in the control group, (9.3 ± 12.3 per participant). The total number, mean and standard deviation ± SD of tracked motor units, per muscle, group are reported in **TABLE 2**.

**TABLE 2.** Total number, mean (± SD) of identified tracked motor units per group and muscle across contractions and conditions.

| Achilles tendinopathy | Mean ± SD | n | Total number |
| --- | --- | --- | --- |
| *Gastrocnemius lateralis* | 9.2 ± 7.1 | 12 | 111 |
| *Gastrocnemius medialis* | 19.8 ± 10.1 | 16 | 317 |
| **Control** |  |  |  |
| *Gastrocnemius lateralis* | 8.0 ± 7.0 | 11 | 88 |
| *Gastrocnemius medialis* | 19.2 ± 13.8 | 14 | 269 |

This table shows mean number of motor units tracked in each muscle across contractions and conditions (*feet-in* and *feet-neutral*), the number of participants (n) who yield motor units and total number of motor units per muscle. standard deviation (SD)

### Motor unit discharge rates

*Gastrocnemius lateralis* motor unit mean discharge rates showed a condition × group interaction (F=5.48, p=0.01, ω2<0.001). Discharge rate was 0.3 (-0.6 to -0.02) pps lower during *feet-in*, compared to *feet-neutral* in the Achilles tendinopathy group and no different, 0.07 (-0.2 to 0.4) pps in the control group (**FIGURE 2A**). Mean discharge rate was not different between groups for *feet-in* 0.77 (-2.4 to 0.9) pps or for *feet-neutral* 0.4 (-2.0 to 1.3) pps. *Gastrocnemius medialis* motor unit mean discharge rate also showed a condition × group interaction (F=71.54 p<0.001, ω2=0.04). Discharge rate was 0.7 (-0.9 to -0.6) pps lower for *feet-in*, in the Achilles tendinopathy group and no different, 0.02 (-0.1 to 0.2) pps in the control group (**FIGURE 2B**). Mean discharge rate was 1.5 (-2.8 to -0.2) pps lower in the Achilles tendinopathy group for *feet-in* compared to controls, and no different between groups for *feet-neutral* 0.7 (-2.0 to 0.5) pps.

**FIGURE 2.**
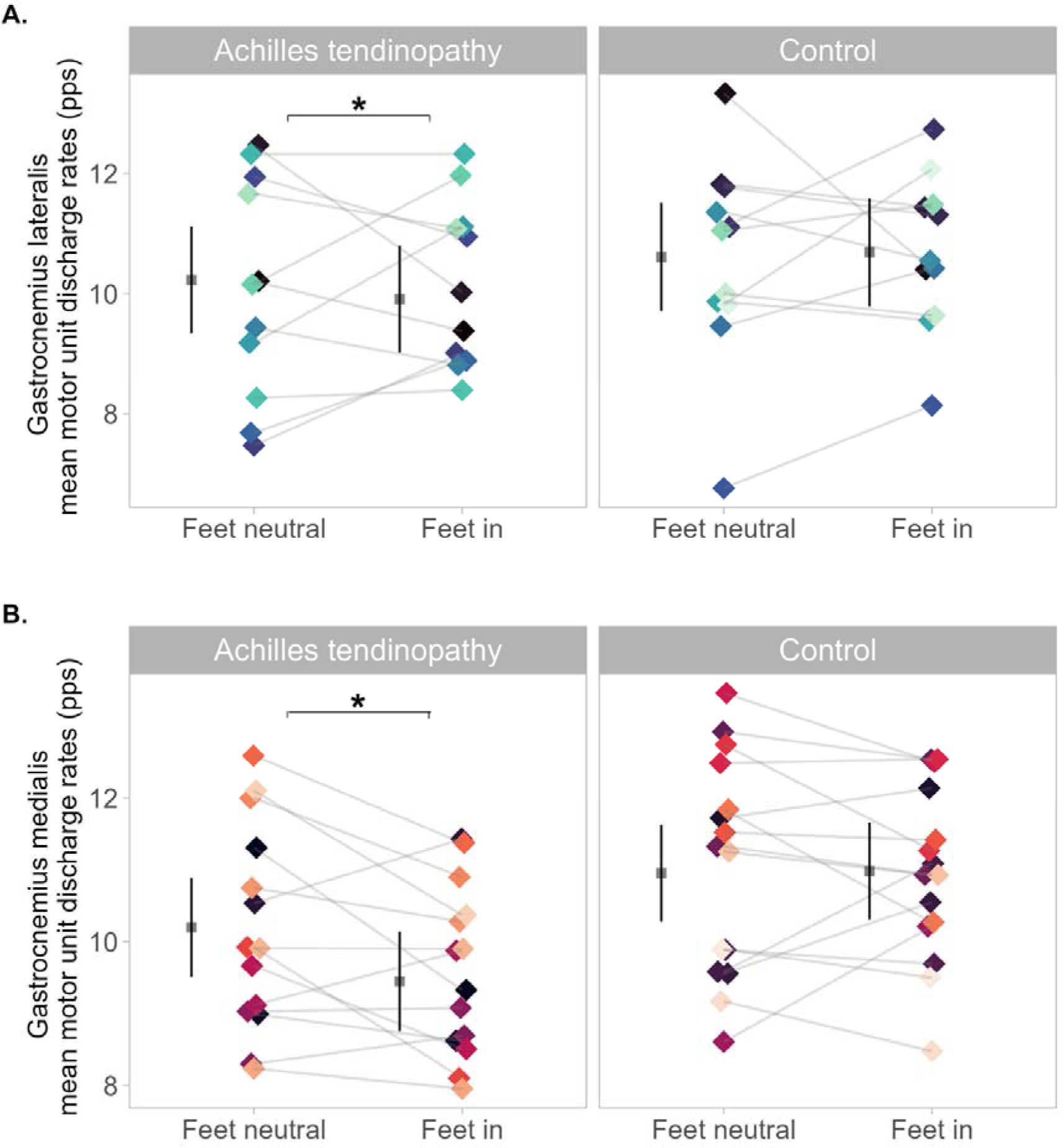
Gastrocnemii motor unit discharge rates with *feet-neutral* and *feet-in* between groups during the tracked motor unit analysis. Each dot represents individual mean motor unit discharge rates across condition, coloured by participants. **A.** represents data from *gastrocnemius lateralis*; and shows lower motor unit discharge rate during *feet-in* (p<0.001) in the Achilles tendinopathy group. **B.** displays data from *gastrocnemius medialis;* and shows lower motor unit discharge rate during *feet-in* (p<0.001) in the Achilles tendinopathy group. Mean and 95% confidence interval are offset to the left to facilitate visualisation. pps = pulse per second. * Denotes statistical difference between conditions.

### Peak isometric torque and single leg heel raise (SLHR) assessments

There were no differences between groups in peak isometric torque (F=2.77, p=0.10), 6.46 (-8.53 to 21.5) N.m. Data for the SLHR was not normally distributed due to the presence of outliers in the control group; Shapiro-Wilk test normality test was p<0.001. After outliers were removed, qqplot and histogram showed a normal distribution and Shapiro-Wilk test normality test was p=0.77. Sensitivity test showed similar statistical results and effect size on both analyses, despite outliers (both results are reported below). The Achilles tendinopathy group performed on average 13.2 (-24.5 to -1.8) fewer SLHR [p=0.024, *d*=-0.77 (-1.5 to - 0.3), large effect size (Lakens et al., 2013)] compared to controls when tested with outliers; and -7.31 (-1.4.9 to -0.7) fewer SLHR [p=0.03, *d*=-0.78 (-1.5 to -0.1)] than controls when outliers were removed (**FIGURE 3**).

**FIGURE 3.**
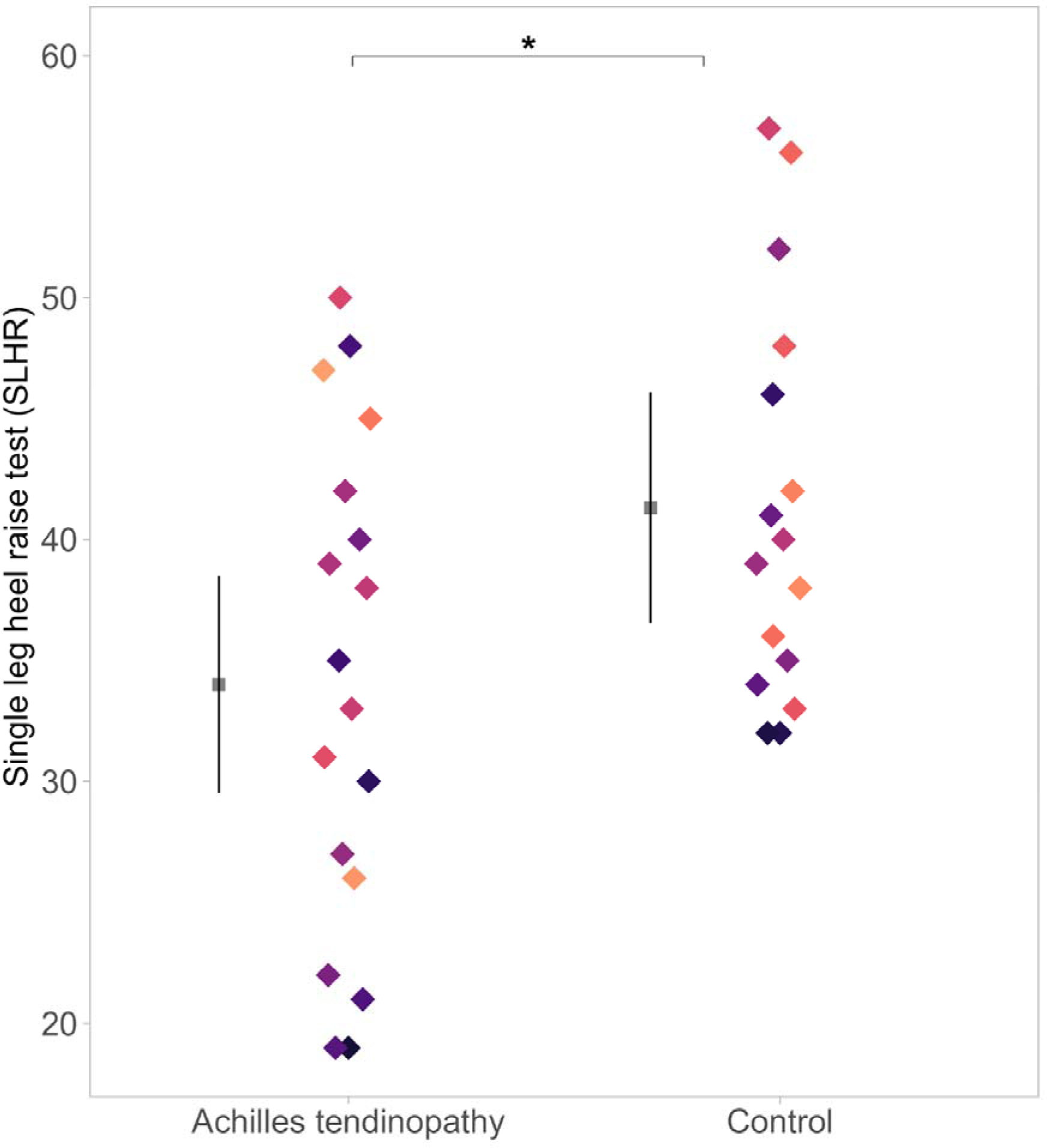
Single leg heel raise (SLHR) test. Mean and 95% confidence interval are offset to the left to facilitate visualisation, data is coloured by participant. Outliers were removed from figure to improve visualisation. * Denotes between-group statistical difference.

## Discussion

This study investigated the effect of different foot position during bilateral isometric standing plantarflexion on the neural drive to the *gastrocnemius lateralis and medialis*, in runners with and without chronic Achilles tendinopathy. Muscle excitation (RMS EMG) and motor unit discharge rates were investigated from *gastrocnemius lateralis* and *gastrocnemius medialis* between conditions (*feet-neutral* and *feet-in*) and compared between groups. We partially confirmed our hypothesis for *gastrocnemius lateralis*, as *feet-in* increased *gastrocnemius lateralis* RMS. However, motor unit discharge rates displayed the opposite response to our hypothesis as it was lower with *feet-in*. Furthermore, our results confirmed the hypothesis for *gastrocnemius medialis*. *Feet-in* reduced RMS and motor unit discharge rates in the Achilles tendinopathy group. Additionally, this study brings new data to support the concept that runners with Achilles tendinopathy have altered neural drive to the *triceps surae,* demonstrated by the reduced motor unit discharge rates to both *gastrocnemii* with *feet-in*. Finally, this study reinforces that plantar flexion endurance deficits are a feature in runners with chronic Achilles tendinopathy, evidenced by poorer SLHR performance.

### Normalised RMS

Our study showed increased normalised RMS during *feet-in* than *feet-neutral* in *gastrocnemius lateralis*, which is in accordance with previous data in the literature. Crouzier et al. (2024) used similar methods, measuring *gastrocnemii* RMS EMG during isometric plantarflexion in healthy individuals and found higher *gastrocnemius lateralis* RMS during *feet-in*. The authors also observed a lower *gastrocnemius medialis* RMS during *feet-in,* compared to *feet out* (feet pointed outwards). Conversely, our data showed lower RMS in *gastrocnemius medialis* RMS during *feet-in* only in the Achilles tendinopathy group, compared to *feet-neutral.* Perhaps, the different position we used, *feet-in* × *feet-neutral* is not sufficient to modify *gastrocnemius medialis* RMS in controls as when comparing *feet-in* × *feet out* as tested by Crouzier and colleagues (Crouzier et al., 2024). Additionally, this helps emphasize that runners with chronic Achilles tendinopathy display different EMG RMS behaviour at different feet position compared to healthy runners. It is important to note that the increase in *gastrocnemius lateralis* RMS observed during the *feet-in* condition was associated to a reduction in *gastrocnemius medialis* in the Achilles tendinopathy group, demonstrating a change in *gastrocnemii recruitment* strategy. This would shift the *gastrocnemius lateralis/medialis* excitation ratio, favouring *gastrocnemius lateralis.* Therefore, this could be an effective strategy to bias *gastrocnemius lateralis* considering the deficits in neural drive observed in individuals with Achilles tendinopathy during various tasks (Crouzier et al., 2020; Mylle et al., 2023).

It should be highlighted that various factors influence sEMG signal amplitudes, such as muscle crosstalk from adjacent deep and superficial muscles; subcutaneous fat thickness; variations in task execution; and the number and type of motor units recruited, and their discharge rates (Lehman & McGill, 1999; Vigotsky et al., 2022). Therefore, greater RMS amplitude does not necessarily translate directly into greater neural drive to the muscle (Vigotsky et al., 2018), thus more specific assessment of individual motor unit discharge behaviour can provide useful information.

### Motor unit discharge rates

We identified motor unit spike trains across contractions and conditions (*feet-in* and *feet-neutral*), which allowed us to gain valuable insight into how each foot position affects each *gastrocnemii* discharge frequencies. *Feet-in* resulted in lower motor unit discharge rates for both *gastrocnemius lateralis* and *gastrocnemius medialis* compared to *feet-neutral*, in the Achilles tendinopathy group. This contradicts our initial hypothesis; rather than increasing motor unit discharge rate, positioning the *feet-in* led to reduced discharge rates in both muscles compared to *feet-neutral*. Furthermore, *gastrocnemius medialis* mean motor unit discharge rates were lower in the Achilles tendinopathy group during *feet-in* than controls. This observation is consistent with our RMS findings, which collectively indicate a reduced neural drive during the *feet-in* condition in the Achilles tendinopathy group. Our *gastrocnemius lateralis* findings from the control group is consistent with another study (Crouzier et al., 2024), which examined various variables, including individual motor unit discharge rates in the *gastrocnemius lateralis* and *medialis* comparing the *feet-in* and *feet out* positions during submaximal isometric plantarflexion in healthy individuals using isokinetic dynamometry. Specifically, there were no differences in *gastrocnemius lateralis* motor unit discharge rates between conditions and lower *gastrocnemius medialis* motor unit discharge rates during *feet-in* condition (Crouzier et al., 2024). A previous study (Hug et al., 2020) found higher motor unit discharge rates in *gastrocnemius lateralis* during *feet-in* and no difference in *gastrocnemius medialis* in non-runners, using a non-tracked motor unit analysis. This contrasts with our results from the *gastrocnemius lateralis* of controls but supports our findings from *gastrocnemius medialis* of the same group. It is important to note that we can only identify a small sample of motor units through decomposition, compared the total number of active motor units contributing to force generation. Therefore, additional motor units may have been recruited but not detected, and this could be speculated as the reason for the opposite findings from EMG RMS. The recruitment of higher threshold larger motor units allows previously recruited active motor units to discharge at a lower frequency while maintaining the same isometric submaximal torque. Additionally, the newly recruited motor units amplify the summation of the global EMG signal, potentially increasing its signal amplitude. Thus, RMS data alone should not be used to inform clinical decision making or guide exercise selection for rehabilitation of individuals with Achilles tendinopathy.

### Neuromuscular measures (SLHR and peak torque)

Isometric peak torque was not different between groups, consistent with findings from other studies (Fernandes et al., 2022; Hasani et al., 2021). This suggests that plantarflexor isometric peak torque is not necessarily lower in runners with chronic Achilles tendinopathy. Conversely, a notable deficit observed consistently across studies, including ours, is in plantarflexor endurance (Fernandes et al., 2021; O’Neill et al., 2019; Quarmby et al., 2023; Silbernagel et al., 2007). The Achilles tendinopathy group performed 18% fewer repetitions to failure compared to controls, aligning with earlier findings from our group (Fernandes et al., 2021). Furthermore, the kinesiophobia scores from the Achilles tendinopathy group represents moderate kinesiophobia (Chimenti et al., 2021). However, our participants still performed a substantial number of SLHRs, without any Achilles tendon pain during testing. These results are similar to values reported in a subgroup of active individuals with Achilles tendinopathy (Hanlon et al., 2021). Therefore, it is unlikely that kinesiophobia is a feature influencing muscle endurance in our sample of runners.

### Strengths and limitations

The main strength of this study was testing the two *gastrocnemius* muscles in one of the most commonly prescribed exercises for of the treatment of runners with Achilles tendinopathy, the standing heel raise, and explored a position variation suggested in the literature to enhance neural drive to *gastrocnemius lateralis*. We were able to track the same motor unit across conditions which provided robust understanding of the neurophysiological mechanisms influencing neural drive influenced by positioning the feet inwards. However, one of the limitations from our study was the ability to record data from up to 64 EMG channels each time. Therefore, we were unable to test both muscles during the same contractions. Additionally, we acknowledge that not testing the *soleus*, a synergist during ankle plantarflexion, was a weakness of this study. Individuals with Achilles tendinopathy display greater contribution from *soleus* over *gastrocnemius lateralis* (Mylle et al., 2023), thus, monitoring *soleus* might have helped explain *gastrocnemius lateralis* reduction in motor unit discharge rates during *feet-in* efforts by the Achilles tendinopathy group. Future research should consider using devices that allow simultaneous recording of multiple channels from all three muscles of the *triceps surae* during the same contractions.

## Conclusion

This study provided novel evidence of the acute effects of foot position during plantarflexion on *gastrocnemius lateralis* and *medialis* RMS and motor unit discharge rates in runners with and without Achilles tendinopathy. While *gastrocnemius lateralis* RMS increased during *feet-in* irrespective of group, motor unit discharge rates reduced for the same condition only in the Achilles tendinopathy group. Additionally, *feet-in* reduced both *gastrocnemius medialis* RMS and motor unit discharge rates in the Achilles tendinopathy group. This suggests that runners with Achilles tendinopathy exhibit an altered *triceps surae* coordination. Therefore, previous results obtained from studies involving a healthy population involving foot position during isometric plantarflexion may not yield the same outcomes in runners with Achilles tendinopathy. Future studies should explore the effect of resistance training protocols utilising different foot positions on individual changes in *triceps surae* motor unit discharge rates to help elucidate associated changes in neural drive during rehabilitation protocols. It is also important to note that runners with Achilles tendinopathy exhibit low *triceps surae* endurance without deficits in peak isometric torque, emphasizing the significance of addressing endurance deficits in rehabilitation for this population.

## Key points

### Findings

These findings demonstrate that adopting a *feet-in* position during plantarflexion can effectively increase *gastrocnemius lateralis* RMS in runners with and without Achilles tendinopathy. Additionally, positioning the feet-inwards, decreases *gastrocnemius medialis* RMS in those with Achilles tendinopathy.

### Implication

Turning the feet inwards shifts the excitation ratio between *gastrocnemius lateralis* and *medialis* in favour of the former, potentially helping to rebalance muscle recruitment in runners with Achilles tendinopathy.

### Caution

While promising, it is uncertain whether performing plantarflexion exercises with feet pointed inwards would provide additional benefits for treating Achilles tendinopathy in runners. Runners with Achilles tendinopathy exhibit an altered coordination strategy within the *triceps surae, compared to those without tendinopathy.* Therefore, strategies for selective *gastrocnemii* recruitment driven by foot position may not function as they do in healthy individuals.

## Data availability

Data and R code are available at https://github.com/GabeFernandess/feet_AT

## Acknowledgments

The authors declare not having received funding for this study.

The authors thank all the volunteers who participated in this study for their contribution to the development and achievement of this research. LBRO is supported by an Alfred Deakin

Postdoctoral Research Fellowship within the Institute for Physical Activity and Nutrition (IPAN) at Deakin University.

## Author contributions

GLF and GST designed the study. GLF conducted the experiments, analysed the data, conducted the statistical analysis, and drafted the first version of the manuscript. All authors critically revised the manuscript and provided valuable feedback. All the authors read and approved this final version of the manuscript.

